# Longitudinal Imaging of Experimental Intracerebral Hemorrhage Pathology with Iodine-enhanced Micro-CT

**DOI:** 10.1101/2025.06.26.661863

**Authors:** Tianjie Zhang, Fan Xia, Mei Fang, Bang Teng, Jiayan Wang, Zexu Wang, Wanting Xia, Dingke Wen, Chuanyuan Tao, Lu Ma, Xin Hu

## Abstract

**Background:** Iodine-enhanced micro-computed tomography (Micro-CT) enables high-resolution three-dimensional imaging of brain architecture. This study aimed to characterize both acute and chronic pathological changes following intracerebral hemorrhage (ICH) using iodine-enhanced micro-CT.

**Method:** Experimental ICH was induced in 8- to 10-week-old C57BL/6 mice (n = 76) via stereotaxic injection of either 0.075 U collagenase IV or 30 μl autologous blood. Iodine-enhanced micro-CT imaging was performed at 4 hours, 1, 3, and 7 days after intracerebral hemorrhage post-ICH to evaluate hematoma formation and erythrolysis. Chronic alterations, including ventriculomegaly and ipsilateral lesion, were assessed at 28 days post-ICH. In parallel, MRI was conducted at 1, 7, and 28 days following autologous blood injection, followed by micro-CT, to facilitate cross-modality quantitative analysis. Lesion volumes were compared between imaging modalities over time.

**Results:** Micro-CT enabled quantification of hematoma volume and erythrolysis in ICH models. Hematomas extended along perivascular pathways toward the cerebral surface in both collagenase- and autologous blood-induced ICH models. At 28 days post-ICH, micro-CT detected ventriculomegaly and hypodense lesions without concurrent expansion of the choroid plexus. Lesion volume measurements derived from micro-CT correlated with those from MRI, enabling quantitatively assessment of iron deposition and hematoma size alterations after ICH.

**Conclusion:** Iodine-enhanced micro-CT provides a robust and high-resolution imaging platform for evaluating hematoma evolution, hemolysis, ventricular enlargement, and chronic brain lesions in experimental ICH. When integrated with MRI, these multimodal imaging approaches enhance the characterization of both hematoma volume and iron deposition following ICH.

## Introduction

Intracerebral hemorrhage (ICH) is the most fatal subtype of stroke and contributes the greatest proportion of disability-adjusted life-years among neurologic disorders^1, 2^. The mass effect and neurotoxicity of hematoma formation causes primary and secondary brain injury after ICH^3^.

Characterization of hematoma evolution in rodent ICH models traditionally relies on histological examinations and magnetic resonance imaging (MRI). Histological methods offer insights into cellular damage and inflammatory response induced by hematoma and erythrolysis^4^. However, they require tissue sectioning, which result in structural distortion and loss of three-dimensional (3D) context^5^. Moreover, histology provides only two-dimensional snapshots, limiting the ability to assess the spatial extent of hemorrhage. MRI enables non-invasive observation of the spatial and temporal hematoma evolution in vivo^6, 7^. T2^*^-weighted imaging is commonly used to estimate hematoma volume in the acute phase and iron deposition in the chronic phase^8, 9^. However, MRI is expensive, has limited spatial resolution, and often produces anisotropic voxels, which constrain its utility for detecting fine structural changes and performing high-precision volumetric analyses^5^.

Micro-CT offers high spatial resolution images and cost-effectiveness compared to MRI^10^, but its ability to visualize soft tissues is inherently limited due to low X-ray attenuation^11^. This limitation can be addressed through the use of contrast agents such as inorganic iodine, which significantly enhances soft tissue contrast^12^. Previous studies have shown iodine-stained mice tissue can effectively identify and quantify the lesion in cerebral cavernous malformations^13^, Alzheimer’s disease^14^ and ischemia stroke^5^.

In this study, we employed ex vivo iodine-enhanced micro-CT imaging to characterize hematoma evolution and associated pathological changes following experimental ICH in mice. Using semi-automated image segmentation and 3D reconstruction, we aimed to: (1) quantify in hematoma volume and erythrolysis ratio at different time points in ICH models; (2) delineate perivascular hematoma morphology during the acute phase; (3) assess changes in the volume of lateral ventricular and choroid plexus during the chronic phase; and (4) compare lesion volume between MRI and micro-CT images across different post-ICH time points.

## Methods

The study was divided into four parts. In the first part, mice received an intracerebral injection of collagenase IV. They were sacrificed at 4 hours, 1 day, 3 days, and 7 days post-injection (n=6 per group), followed by micro-CT imaging. In the second part, mice were injected with 30ul autologous blood, and euthanized at 4 hours, 1 day, 3 days, 7 days post-injection (n=6 per group), with subsequent micro-CT scans. In the third part, mice were randomized to receive intracerebral injections of 30ul autologous blood, 0.075 U collagenase type IV, or sham operation (n=6 per group). All groups were euthanized for 28 days post-injection, and then micro-CT scans were performed. In the fourth part, mice had an injection of 30ul autologous blood, and underwent MRI at 1 day, 7 days, and 28 days post-injection (n=4 per timepoint). Then mice were then euthanized for micro-CT. A total of 76 C57Bl/6 male mice (8–10 weeks, 20–25 g) were used, 5 mice died after modeling and were excluded from this study.

### Animal preparation

All animal procedures and experimental protocols followed the Ethical Principles in Animal Experimentation established by Sichuan University and were approved by the Animal Care and Use Ethics Committee of Sichuan University (Ethics Approval Number: 20240509005). The mice were supplied by GemPharmatech Co., Ltd. and housed in a controlled environment maintained at a temperature of 23–25 °C and a humidity level of 45–50%. They were kept on a 12-hour light-dark cycle with unlimited access to food and water.

### Collagenase IV and autologous blood injection

Intracerebral injection was conducted as previously described^15^. Briefly, mice were anesthetized with isoflurane and secured in a stereotaxic frame. A 0.7 mm cranial burr hole was drilled. For collagenase IV induced ICH model, a 10 μL syringe was used to inject 0.6 μL of saline containing 0.075 U collagenase IV (Sigma-Aldrich) into the striatum (coordinates: 0.2 mm anterior, 3.6 mm ventral and 2.2 mm lateral to the bregma) at a rate of 0.12 μL/min. For autologous blood induced ICH model, mice were anesthetized with isoflurane, and the right femoral artery was cannulated to obtain autologous blood. A 26-gauge needle was inserted stereotaxically into the right striatum (coordinates identical to above). Blood (30 ul) was infused into the striatum at a rate of 3 ul/min. Mice in sham group underwent needle insertion without infusion. Following the injection, the needle was retained for 5 minutes. The burr hole was sealed with bone wax, and the scalp incision was closed with sutures.

### Iodine staining and Histology

Following isoflurane-induced anesthesia, mice underwent transcardially perfused with 4% paraformaldehyde (PFA). Subsequently, brains were extracted and subjected to post-fixation in 4% PFA at 4°C for a minimum of 5 days. Dehydration was then performed via a sequential ethanol gradient (30%, 50%, 70%, 80%, 90%, each for 2 hours). Tissues were subsequently stained for a duration of 24 hours in 90% methanol containing 1% iodine at room temperature. After iodine staining, the brains underwent rehydration using 70% and 30% ethanol solutions (1 hour each) at room temperature. Samples were then wrapped in Parafilm® to minimize dehydration, and subjected to micro-CT imaging. For short-term preservation, brains were stored at 4°C.

After imaging, brains were then embedded in optimal cutting temperature (O.C.T.) compound, and coronal sections (18μm thickness) were obtained using a cryostat. Erythrolysis within the hematoma was histologically confirmed by hematoxylin and eosin (H&E) staining, in accordance with established protocol^16^.

### Micro-CT imaging and reconstruction

Iodine-stained mouse brains underwent scanning via a BRUKER micro-CT scanner (SkyScan1276, BRUKER). The operational parameters employed included a voltage of 60 kV and a current of 200 µA. A spatial resolution of 16 µm was achieved, with a rotational increment of 0.2° across a complete 360° revolution, resulting in 1801 projections. A 0.5 mm aluminum filter, combined with a frame averaging value of 2, was utilized. The duration for scanning each brain was approximately 8 minutes. Reconstruction of the acquired images was performed using the NRecon software (BRUKER). While normalization settings were applied uniformly across all brains, fine-tuning was implemented individually to optimize reconstruction quality for each sample.

### Semi-automatic segmentation and quantification

The reconstructed micro-CT files were transformed into. ims format using Imaris software. Surface rendering was then conducted, employing a surface detail level of 16. Machine learning segmentation was adopted following procedure: The slicer extent parameter was configured to a value of 16, followed by manual annotation of both target regions and background areas. Following the preliminary training and prediction, the segmentation outcomes underwent visual assessment. The selections for foreground and background were iteratively refined through repetitive training cycles until an acceptable level of segmentation precision was attained. For quantitative evaluation, volumes were derived from the detailed plate within the software’s statistics module.

### Magnetic resonance imaging

Mice were anesthetized with a 2% isoflurane/air mixture and body temperatures were maintained using a forced-air heating system.

T2- and T2^*^-weighted images (T2^*^WI, T2WI) were acquired at days 1, 7, 28 after autologous blood injection using a 7.0-T Bruker Biospec USR 70/30 MRI system as described previously^17^. Parameters for T2-weighted images: field of view (FOV) = 20 × 20 mm, repetition time (TR) = 2519.22ms, echo time (TE) = 33ms, acquisition matrix = 256 × 128, slice thickness = 0.5mm. T2^*^-weighted images were acquired with the following parameters: FOV = 20 × 20 mm, TR= 150 ms, TE = 6 ms, acquisition matrix = 256 × 256, slice thickness = 0.5 mm. The T2^*^ lesion volume was measured over the total areas in all slices by multiplying the section thickness.

### Comparison of micro-CT and MRI results

Due to tissue shrinkage during ethanol dehydration of micro-CT sample preparation, the proportion of hematoma in the brain obtained from micro-CT imaging were calculated to facilitate comparison with MRI results.

First, identify the lesion level on T2^*^ images. Using the previously described method, calculate both the T2^*^ lesion volume and the brain volume at the corresponding level. The T2^*^ lesion ratio was calculated by dividing the T2^*^ lesion volume by the corresponding brain volume.

Next, on micro-CT images, select the anatomically corresponding level matching the MRI lesion level. Calculate both the hematoma volume and the micro-CT brain volume corresponding to MRI lesion level, thereby obtaining the micro-CT hematoma ratio by dividing the hematoma volume by the micro-CT brain volume.

Since T2^*^ *sequences simultaneously detect both hematoma and iron deposits, while micro-CT is only sensitive to hematoma*, the hematoma proportion is calculated as the ratio of micro-CT hematoma ratio to T2^*^ lesion ratio. The iron proportion was subsequently computed by subtracting the hematoma proportion from 100%.

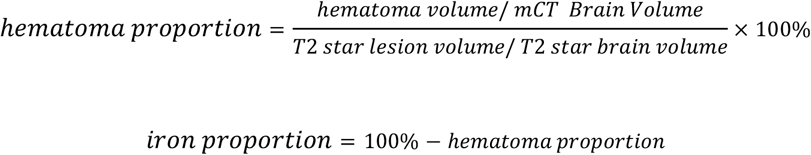

### Statistical analysis

Data analysis was conducted utilizing GraphPad Prism software (Version 9.0). Experimental results were quantified and expressed as mean ± standard deviation (SD). Group comparisons were evaluated through unpaired student t-test and one-way ANOVA with Tukey’s multiple comparisons test (^**^p < 0.01, ^*^p < 0.05).

## Results

### Micro-CT Assessment of Hematoma Volume in a Collagenase-Induced Intracerebral Hemorrhage Model

To evaluate hematoma evolution, mice subjected to intrastriatal injection of collagenase IV were euthanized at 4 hours, 1 day, 3 days, and 7 days post-ICH. Brains were harvested for ex vivo iodine staining and micro-CT imaging (Figure 1A). Three-dimensional reconstruction of hyperdense hematoma regions was performed using Imaris software. Representative reconstructed images revealed a porous morphology characteristic of the collagenase-induced hematomas (Figure 1B-C). Quantitative analysis demonstrated a significant increase in hematoma volume at 1 day compared to 4 hours post-injection (3.09±0.45 mm^3^ vs 1.98±0.42 mm^3^, p<0.01), indicating hematoma expansion during this interval (Figure 1D). Hematoma volume subsequently declined significantly by day 3 (vs 0.99±0.45 mm^3^, p<0.01) and was nearly resolved by day 7 (vs 0.18±0.17 mm^3^, p<0.01).

**Fig. 1.**
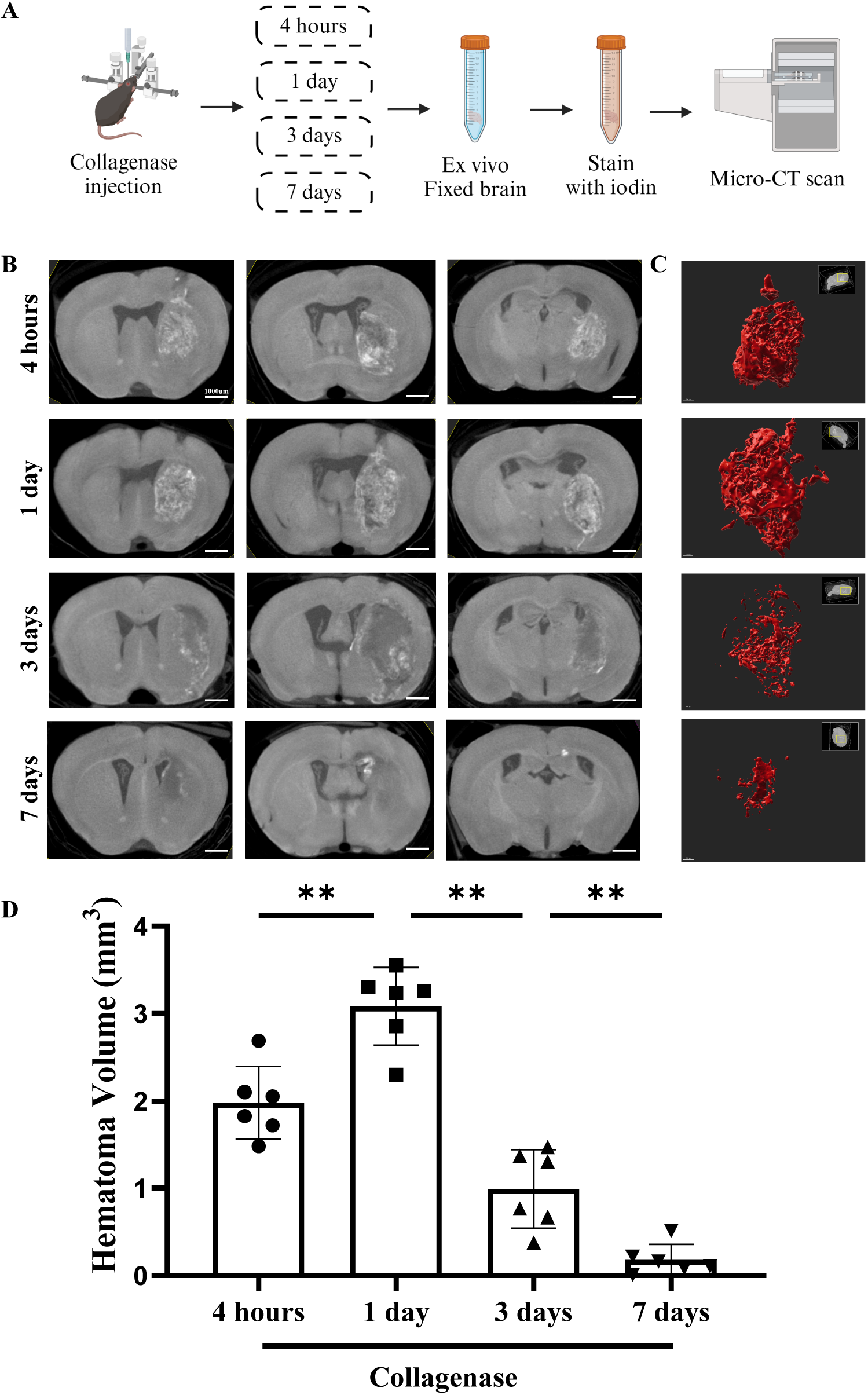
Hematoma volume of collagenase-induced intracerebral hemorrhage model by Micro-CT imaging. (A) Experimental workflow for micro-CT scanning in a collagenase-induced ICH model. Time points of analysis include 4 h, 1 d, 3 d, and 7 d post-injection. Representative micro-CT images (B) and three-dimensional reconstructions (C) showed hematoma morphology at specified time points. (D) Quantitative data of hematoma volume at 4 h, 1 d, 3 d, and 7 d post-injection (n = 6 per timepoint). Data are presented as mean ± SD. (^**^P < 0.01).

### Micro-CT Assessment of Hematoma Volume in an Autologous Blood-Induced Intracerebral Hemorrhage Model

In mice receiving autologous blood injections, brains were collected at the same time points for ex vivo iodine staining and micro-CT imaging (Figure 2A). Representative images demonstrated solid, non-porous hematomas in the autologous blood model, morphologically distinct from those in the collagenase model (Figures 2B-C). Quantitative analysis revealed no significant difference in hematoma volume between the 4 hours and 1 day post-injection (4.16±0.61 mm^3^ vs 3.82±0.95 mm^3^, p>0.05, Figure 2D). Hematoma volume began to decrease by day 3 (vs 2.67±0.69 mm^3^, p<0.05), with a significant reduction by day 7 (vs 0.50±0.21 mm^3^, p<0.01).

**Fig. 2.**
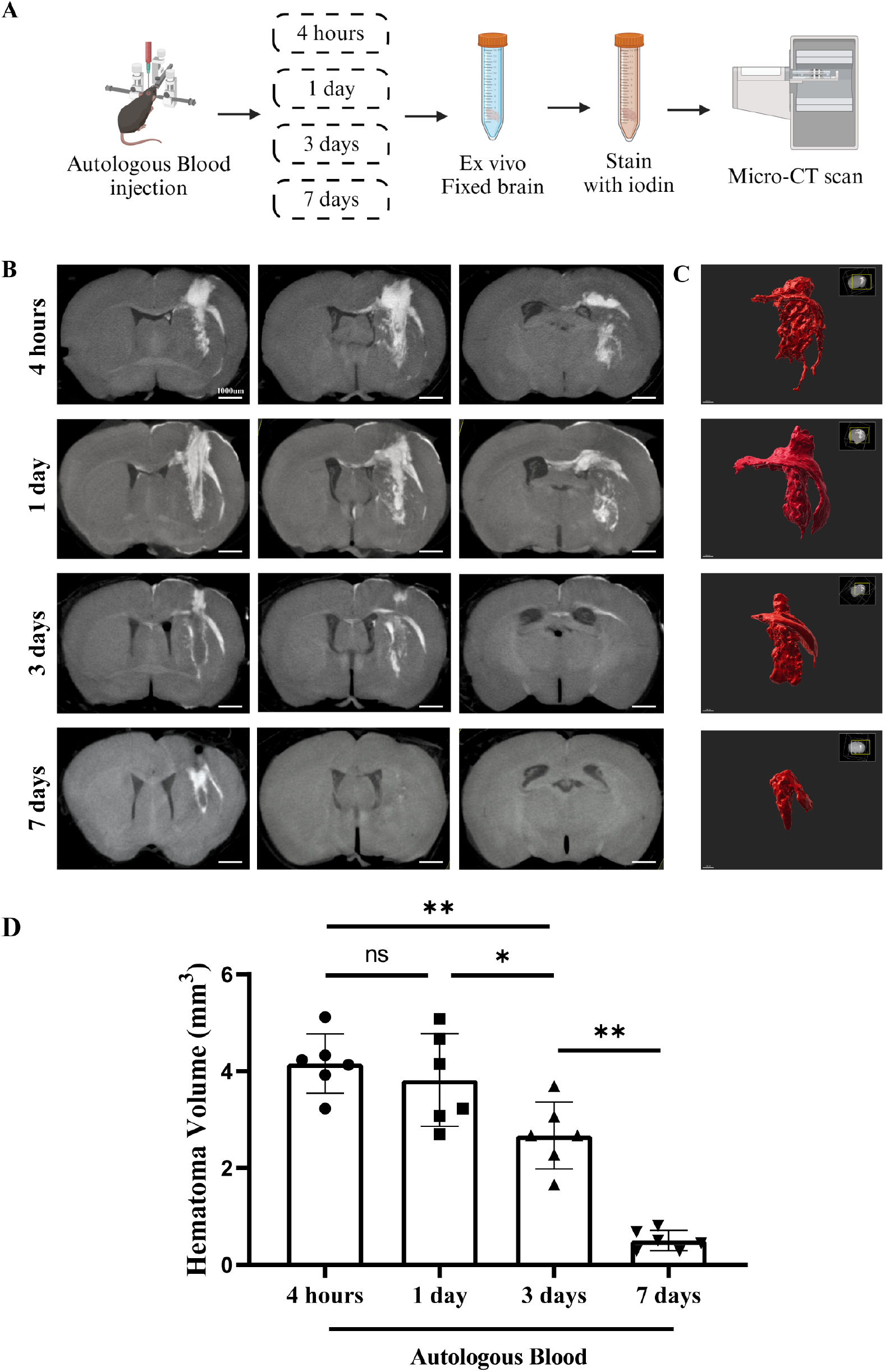
Hematoma volume of autologous blood-induced intracerebral hemorrhage model by Micro-CT imaging. (A) Experimental workflow for micro-CT scanning in a autologous blood-induced ICH model. Scans were conducted at 4 h, 1 d, 3 d, and 7 d following autologous blood injection. Representative micro-CT images (B) and three-dimensional reconstructions (C) demonstrated hematoma morphology at indicated timepoints. (D) Quantification of hematoma volume at 4 h, 1 d, 3 d, and 7 d post-injection (n = 6 per timepoint). (^*^P < 0.05, ^**^P < 0.01).

### Micro-CT Imaging Enables Visualization and Quantification of erythrolysis after Intracerebral Hemorrhage

In the autologous blood model, micro-CT imaging revealed hypodense regions embedded within hyperdense hematomas. These regions were presumed to represent erythrolysis, in line with prior studies using non-hypointense signal within iron signal on T2^*^-weighted MRI^16, 18^. Histological validation confirmed that hypodense areas corresponded to zones of erythrolysis (Figure 3A-B). To quantify erythrolysis progression after ICH, three-dimensional reconstructions of hypodense area and hyperdense hematoma regions were generated (Figure 3C), and the ratio (erythrolysis volume/total hematoma volume x100%) was calculated. At 4 hours post-ICH, erythrolysis was minimal (0.76±0.34%, Figure3D). By day 1, the ratio increased significantly (vs 4.15±1.41%, p<0.05), peaked at day 3 (vs 12.50±2.95%, p<0.01), and then declined slightly by day 7 (vs 10.72±1.96%, p>0.05).

**Fig. 3.**
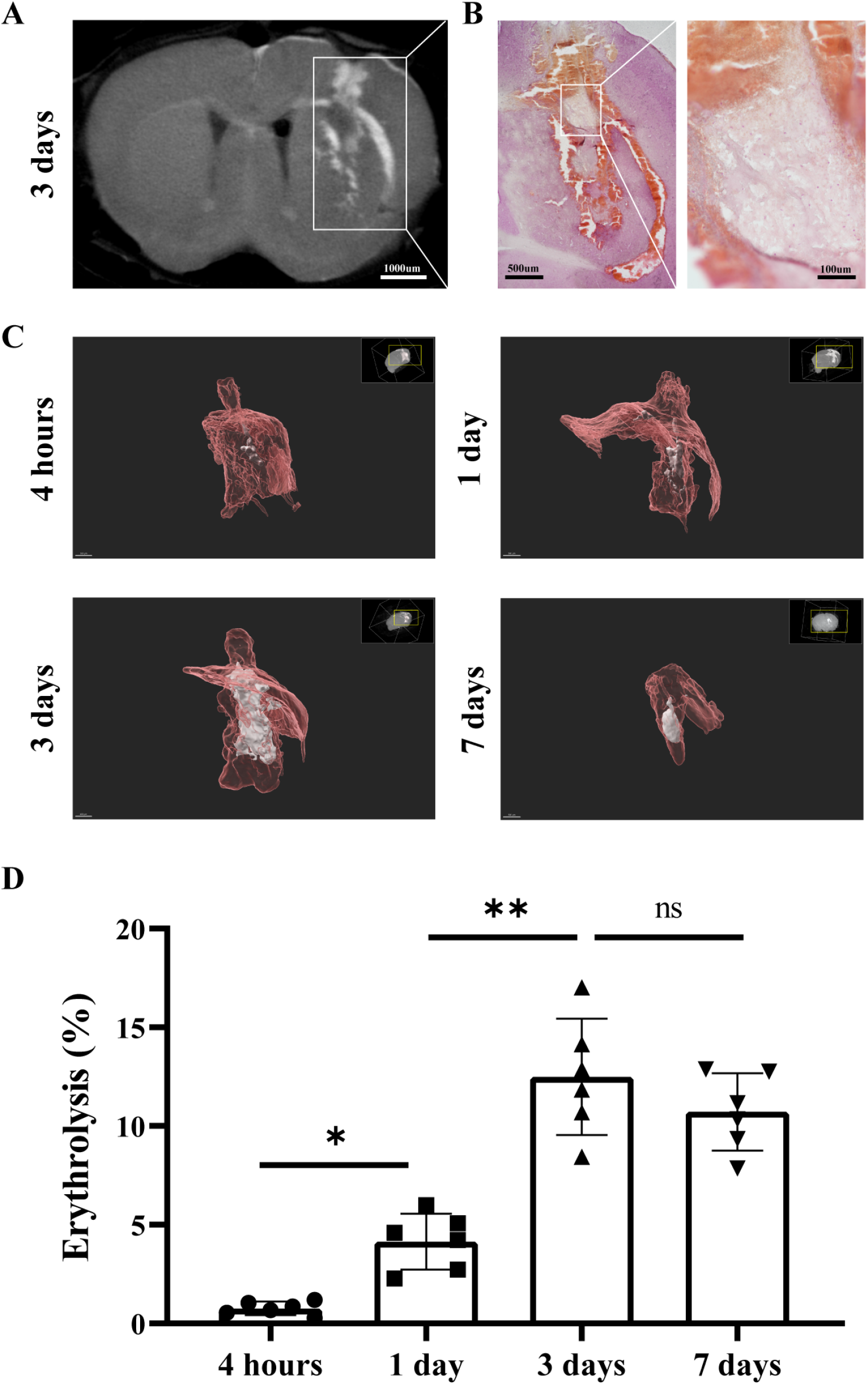
Micro-CT characterization of hemolysis following intracerebral hemorrhage. (A) Coronal micro-CT image at 3 d after autologous blood induced hemorrhage. White box highlights hypodense regions within the hyperdense hematoma. (B) Histological validation by H&E staining of the corresponding section. Boxed area demonstrates erythrolysis zones matching hypodense regions in micro-CT images. (C) Three-dimensional reconstruction images of hematoma and hemolysis 4 hours, 1 day, 3 days and 7 days post injection. (D) Quantification of hemolysis ratio at 4 h, 1 d, 3 d, and 7 d post-injection (n = 6 per timepoint). (^*^P < 0.05, ^**^P < 0.01).

### Hematoma Extends Along Perivascular Pathways to the Brain Surface after Intracerebral Hemorrhage

Prior studies suggested that intracerebral hematomas spread along perivascular pathways after ICH^19^. In our study, micro-CT imaging revealed hematoma extend along perivascular pathways to the surface of the brain in both collagenase- (Figure 4A) and autologous blood- (Figure 4B) induced ICH models. Quantitative analysis showed a significant increase in the number of hematomas reaching the brain surface in the collagenase model from 4 hours to day 1 (1.83±1.47 vs 4.14±1.07, p<0.05, Figure 4C), which may due to persistent bleeding after collagenase injection. In contrast, no significant difference was observed between 4 hours and 1 day in the autologous blood model (6.00±1.41 vs 5.83±1.72, p>0.05). Scatter plots showed a positive correlation between hematoma volume and the number of hematomas reaching the brain surface. Scatter plot analysis revealed a positive correlation between hematoma volume and the number of hematomas reaching the brain surface at 4 hours and 1 day in the autologous blood and collagenase model (r=0.79, p<0.01, Figure 4D), suggesting that hematoma volume is a critical factor influencing perivascular hematoma extension.

**Fig. 4.**
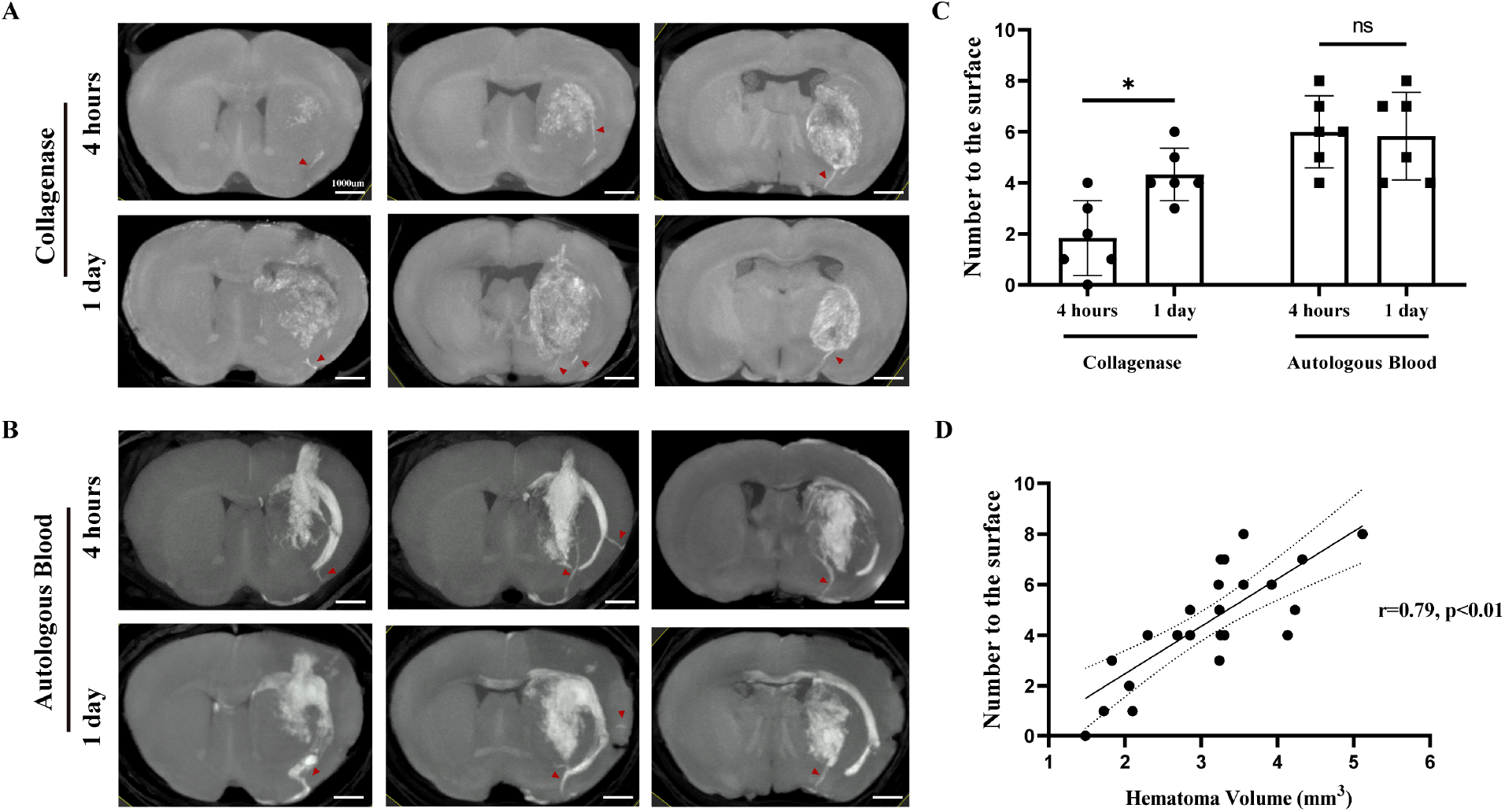
Micro-CT visualization of hematoma extended along perivascular pathways to the brain surface. Representative micro-CT images of the autologous blood (A) and collagenase (B) injection model at 4 h and 1 d post-induction (n = 6 per group). Red arrows showed hematoma distribution along perivascular routes. **(C)**, Quantification of the number of perivascular hematomas reaching the brain surface. (D) Scatter plot analysis of hematoma volume and the number of hematomas reaching the brain surface at 4 hours and 1 day post-induction in the autologous blood and collagenase model revealed a positive correlation between these two parameters. (^*^P < 0.05, ^**^P < 0.01)

### Ventriculomegaly and Ipsilateral Hypodense Lesions in Chronic Phase after Intracerebral Hemorrhage

At 28 days post-ICH, ex vivo micro-CT imaging was performed to assess chronic structural changes (Figure 5A). Three-dimensional reconstructions showed significant lateral ventricle enlargement in both the collagenase (3.55±0.59 mm^3^ vs 1.55±0.29 mm^3^, p<0.01) and autologous blood (vs 2.13±0.23 mm^3^, p<0.05) models (Figures 5B-C). Meanwhile, the volume of lateral ventricle in collagenase model was larger than that in autologous blood model (p<0.05). There were no significant differences in choroid plexus volume between the autologous blood group and the sham group (0.23 ±0.02 mm^3^ vs 0.24±0.02 mm^3^, p>0.05, Figure 5D), nor between the collagenase group and the sham group (vs 0.20±0.01 mm^3^, p>0.05, Figure 5D).

**Fig. 5.**
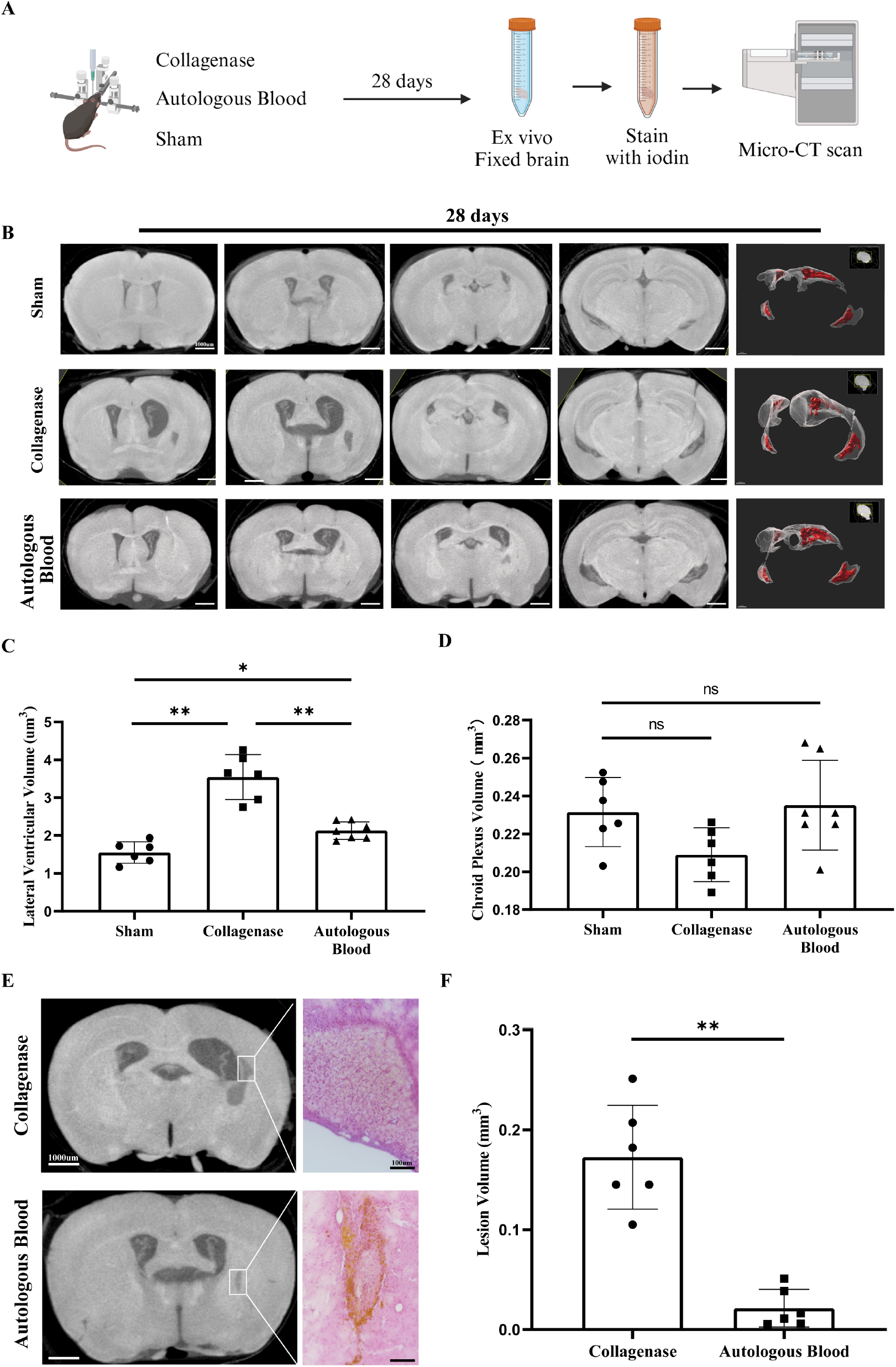
Chronic lesions in experimental intracerebral hemorrhage models revealed by micro-CT imaging. (A) Experimental workflow for micro-CT evaluation in chronic phase of ICH models (B) Representative micro-CT images and three-dimensional reconstructions of lateral ventricular (transparent) and choroid plexus (red) at 28 d post-intervention in autologous blood injection, collagenase-induced hemorrhage, and sham-operated groups (n = 6 per group). Quantitative analysis of lateral ventricular volume(C) and choroid plexus (D) in collagenase, autologous blood and sham groups. (C) Low-signal regions on micro-CT images and corresponding HE-stained sections at 28 d after autologous blood and collagenase injection. (F) Quantitative analysis of chronic lesion volume on micro-CT images 28 d after injection.

Hypodense regions in the ipsilateral hemisphere were observed in both models. Histological analysis of micro-CT–scanned brains revealed that in the collagenase model, these regions were predominantly composed of nucleated cells with minimal hemosiderin deposition. In contrast, the autologous blood model displayed evident hemosiderin accumulation surrounding fibrotic tissue (Figure 5E).

Three-dimensional reconstruction and volumetric quantification of the hypodense areas revealed that the collagenase group exhibited significantly larger hypodense volumes compared to the autologous blood group (Figure 5F), indicating more extensive chronic tissue damage.

### Dynamic Evaluation of Hematoma Size and Iron Content with Micro-CT and MRI

To evaluate the hematoma size and iron deposition after experimental ICH, mice injected with autologous blood underwent *in vivo* MRI scans at 1-, 7-, and 28-days post-injection (*n* = 6 per timepoint), followed by ex vivo micro-CT imaging (Figure 6A).

**Fig. 6.**
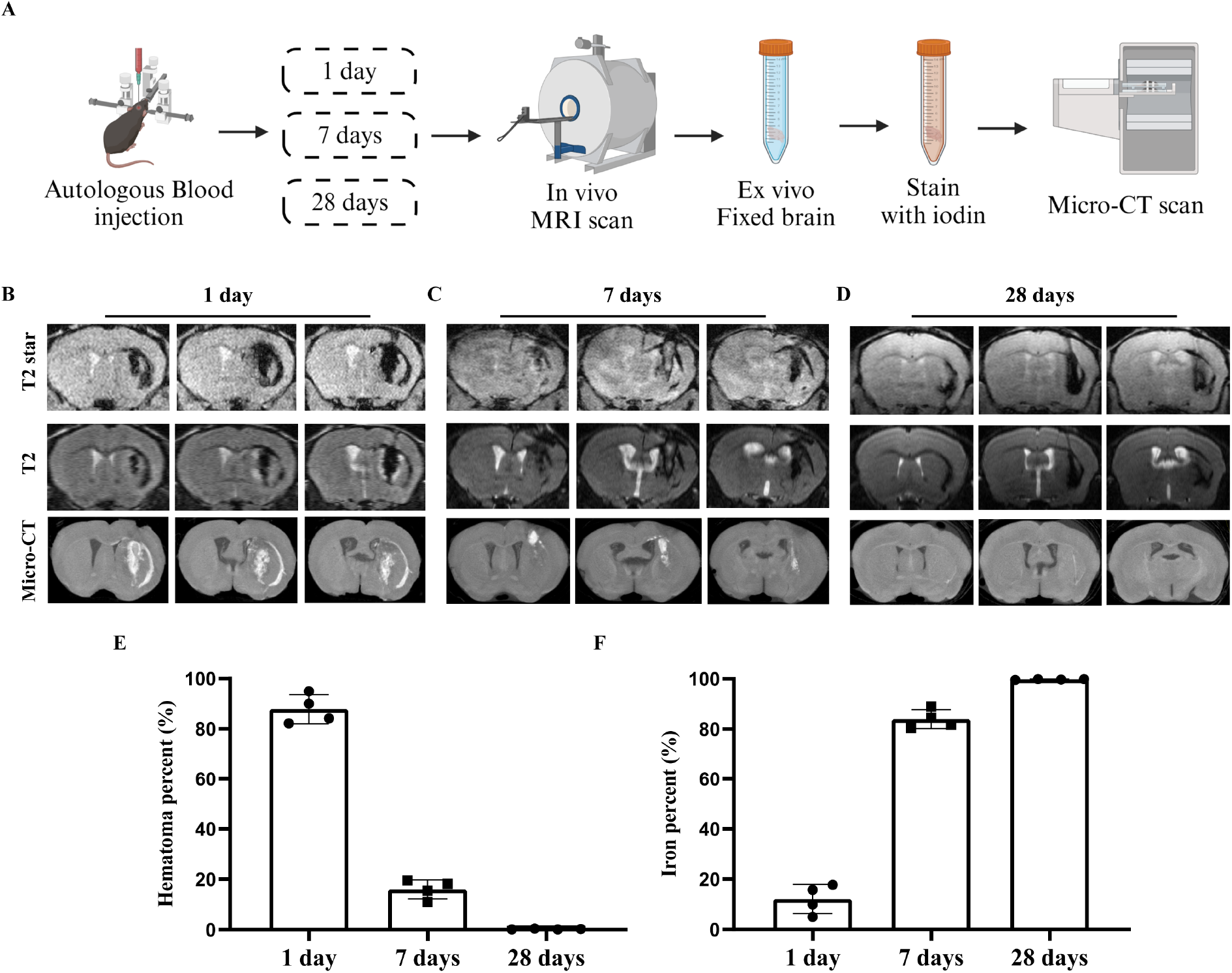
Multimodal imaging comparison of experimental intracerebral hemorrhage. (A) Experimental workflow for multimodal imaging comparison of autologous blood induced intracerebral hemorrhage at 1 d, 7 d, and 28 d post-injection. Representative T2^*^ (top), T2 (middle) and micro-CT (bottom) images at 1 d (B), 7 d (C), and 28 d (D) after injection. Quantitative data of hematoma percent and iron deposition percent at 1d, 7d, and 28d after autologous blood injection. (^*^P < 0.05, ^**^P < 0.01).

At day 1, micro-CT and T2^*^-weighted MRI showed similar lesion patterns (Figure 6B). The quantification data showed hematoma proportion was 87.86 ± 5.79%, while the iron proportion was 12.14 ± 5.79% (Figure6 E). By day 7 post-injection, the T2^*^ lesion area was significantly larger than the hematoma region detected by micro-CT (Figure 6C). The hematoma proportion showed a significant decrease (vs. 16.07 ± 3.75%, p < 0.01, Figure 6D), whereas the iron deposition proportion increased compared to day 1(vs. 83.93 ± 3.75%, p< 0.01 Figure 6 E). At 28 days post-injection, the representative micro-CT images showed almost no hyperdense area (Figure6 D), the hematoma was nearly completely cleared (vs. 0.24 ± 0.15%, p<0.01), while T2^*^-weighted imaging persistently demonstrated prominent iron deposition (99.76 ± 0.15%, p < 0.01).

## Discussion

The main findings of this study are summarized as follows: (1) Micro-CT imaging enables quantitative evaluation of hematoma volume and erythrolysis in experimental ICH. (2) intracerebral hematoma extends along perivascular routes to the cerebral surface in both collagenase and autologous blood models. (3) significant ventriculomegaly was observed without choroid plexus volume increasing following chronic phase after ICH. (4) the combined use of micro-CT and MRI facilitates characterization of the temporal dynamics of hematoma resolution and iron deposition following ICH.

In the acute phase of ICH, previous studies have have demonstrated that hematoma size can be assessed using T2^*^-weighted MRI, with non-hypointense regions within hypointense hematomas indicative of erythrolysis^4, 16, 20^. In chronic phase, T2^*^-weighted signal predominantly reflect hemosiderin-laden iron deposition rather than intact hematoma^9^. However, it is difficult to distinguish hematoma from iron by only MRI, especially in the subacute phase after ICH. In this study, we employed iodine-enhanced micro-CT to quantitatively assess both hematoma volume and erythrolysis. When combined with MRI, this dual-modality approach offers a powerful framework for delineating the dynamic evolution of hematoma and iron over time.

Our study also provides imaging-based evidence for hematoma extension along perivascular pathways. Hematoma spread along the perivascular spaces has been previously described in both preclinical and clinical contexts^19, 21, 22^. We visualized this phenomenon using high-resolution micro-CT in both collagenase- and autologous blood-induced ICH models. These findings offer additional insight into the anatomical dissemination of blood components within the brain parenchyma and may have implications for understanding hematoma expansion and clearance.

Ventricular enlargement is a well-recognized marker of brain atrophy during chronic phase after ICH^18^. Concurrently, increases in choroid plexus volume have been associated with inflammatory responses in other neuropathological contexts ^23^. In this study, we used micro-CT to reconstruct and quantify lateral ventricular and choroid plexus volumes 28 days after ICH. Our data showed the lateral ventricular enlargement accompanied by a nonsignificant reduction in choroid plexus volume during the chronic phase in both collagenase-induced and autologous blood-induced ICH models. These observations imply that post-ICH ventriculomegaly may be more closely attributable to brain atrophy rather than choroid plexus inflammation at chronic phase of ICH. Future studies should further explore the relationship between choroid plexus volume and hydrocephalus in intraventricular (IVH) and subarachnoid hemorrhage (SAH) models, where hydrocephalus is more pronounced.

This study has several limitations. First, ex vivo iodine staining requires brain dehydration, which results in brain volume shrinkage. Although three-dimensional reconstruction showed no significant differences in brain volume across experimental groups following dehydration and iodine staining, direct volumetric comparisons between ex vivo micro-CT and in vivo MRI remain limited due to tissue shrinkage during the staining process.

In conclusion, this study demonstrates that iodine-enhanced micro-CT is a robust high-resolution modality for quantifying hematoma volume, erythrolysis ratio, ventricular enlargement, and chronic lesion burden in experimental ICH. The integration of MRI and micro-CT imaging provides valuable insights into temporal changes of hematoma volume and iron deposition following ICH. Future investigations should explore the application of iodine-enhanced micro-CT in other hemorrhagic stroke subtypes, including subarachnoid hemorrhage or intraventricular hemorrhage, to enhance its translational relevance.

## Data sharing statement

The data used or analyzed in the current study are available from the corresponding author upon reasonable request.

## Acknowledgements

This work was supported by the National Natural Science Foundation of China (82201450 and 82201453), the Natural Science Foundation of Sichuan Province (2023NSFSC1560), and the Sichuan Science and Technology Program (2023NSFSC1557).

